# Gentamicin induction of the gonococcal *hicAB* toxin-antitoxin encoding system and impact on gene expression influencing biofilm formation and in vivo fitness in a strain-specific manner

**DOI:** 10.1101/2025.06.11.659166

**Authors:** Concerta L. Holley, Vijaya Dhulipala, Adriana Le Van, Jacqueline T Balthazar, Vincent J. Oliver, Ann E. Jerse, William M. Shafer

**Affiliations:** Department of Microbiology and Immunology, Emory University School of Medicine, Atlanta, Georgia, USA; Department of Microbiology and Immunology, Uniformed Services University, Bethesda, Maryland, USA; The Emory Antibiotic Resistance Center, Emory University School of Medicine, Atlanta, Georgia, USA; Laboratories of Bacterial Pathogenesis, Veterans Affairs Medical Center, Decatur, Georgia, USA

**Keywords:** *Neisseria gonorrhoeae*, gentamicin, toxin-antitoxin, biofilm, stress response

## Abstract

The continued emergence of *Neisseria gonorrhoeae* (Ng) isolates resistant to first-line antibiotics has focused efforts on understanding how alternative therapies such as expanded use of gentamicin (Gen) might counteract this global public health problem. Focusing on Gen as a viable alternative antibiotic for treatment of gonorrheal infections, we used RNA-Seq to determine if sub-lethal levels of Gen might impact gonococci on a transcriptional level. We found that sub-lethal Gen levels altered expression of 23 genes in Ng strain FA19. Many of the differentially regulated genes were associated with known stress responses elaborated by Ng under different harmful conditions. We found that the transcripts of the *hicAB* operon, which encodes a putative HicA-HicB toxin-antitoxin system that is encoded by tandem genes with the prophage Ngo φ3, were increased in response to Gen. While loss of *hicAB* did not impact gonococcal susceptibility to a variety of antimicrobial agents or harmful environmental conditions it did reduce biofilm formation in Ng strains F62, FA1090, WHO X and CDC200 but not that of strain FA19. Further, in strain F62, but not FA19, loss of *hicAB* reduced the *in vivo* fitness of Ng during experimental lower genital tract infection of female mice. Further, we found that expression of *hicAB* can influence levels of the *norB* transcript, which encodes the nitrate reductase shown previously to be upregulated in gonococcal biofilms. We propose that sub-lethal Gen has the capacity to influence gonococcal pathogenesis through the action of the HicAB toxin-antitoxin system.

**Importance:** During antibiotic treatment bacteria can be exposed to sub-lethal levels that could serve as a stress signal resulting in changes in gene expression. The continued emergence of multi-drug resistant strains of Ng has rekindled interest in expanded use of gentamicin (Gen) for treatment of gonorrheal infections. We report that sub-lethal levels of Gen can influence levels of Ng transcripts including that of the gonococcal *hicAB*-encoded toxin-antitoxin (TA) locus, which is embedded within an integrated prophage, While loss of this TA locus did not impact Ng susceptibility to Gen it reduced the biofilm forming ability of 4/5 Ng strains. Further, in an examined strain in this group we found that Ng fitness during experimental infection was negatively impacted. We propose that that levels of the *hicA-hicB* transcripts can be increased by sub-lethal levels of an antibiotic used in treatment of gonorrhea and that this could influence pathogenicity.

## Introduction

*Neisseria gonorrhoeae* (Ng) is the obligate human pathogen that causes gonorrhea, the second most commonly reported sexually transmitted bacterial infection with an estimated 87 million cases worldwide per year (1). Since the start of modern chemotherapy in the late 1930s Ng has displayed the remarkable ability to express clinical resistance to all introduced antibiotics (2) and multi-drug resistant (MDR) isolates have been reported globally in recent years (3–5). Increasing antibiotic resistance of Ng is a growing global concern and Ng is considered an urgent public health threat pathogen by both the Centers for Disease Control and Prevention (CDC) and the World Health Organization (WHO) (2, 4, 6).

The continued emergence of Ng isolates resistant to first-line antibiotics has re-invigorated interest in alternative antibiotic therapies such as expanded use of gentamicin (Gen). This aminoglycoside is known to have anti-gonococcal action and has extensive use in many countries when recommended first-line gonorrheal therapy failed (5) and in Malawi as a front-line empirical gonorrheal therapy in combination with doxycycline (7, 8). We hypothesized that expanded use of Gen may promote emergence of Ng with clinically relevant resistance and conducted earlier studies to understand how decreased susceptibility of Ng to Gen might develop (9). In this regard, we found that a single amino acid change in *fusA*, which encodes elongation factor G (EF-G) could significantly decrease Ng susceptibility to Gen (9, 10).

The majority of verified treatment failures are associated with pharyngeal infections (11–13), and despite Gen being recommended as alternate therapy for treatment of urogenital gonorrhea, it cannot reliably alone eradicate Ng from the pharynx (14), which is likely due to pharmacokinetic (PK) properties of Gen that impact its tissue penetration ability (15). Pharmacodynamic (PD) studies have shown killing of Ng with Gen concentrations above the minimum inhibitory concentration (MIC) within two to three hours *in vitro* (16). However, a study in beagles demonstrated that Gen has less than a two-hour half-life *in vivo* and reaches a suboptimal plasma concentration within an hour in mice (17, 18). Thus, in patients treated with Gen to cure a gonorrheal infection, the surviving Ng, especially those residing in the pharynx, may be exposed to sub-lethal Gen.

Understanding how Ng responds to sub-lethal levels of Gen has implications for advancing knowledge regarding the possible impact of antibiotic stress on bacterial responses that might influence pathogenicity. In relation to our present study, an earlier report by Hoffmann *et. al.* showed that sub-lethal levels of aminoglycosides, especially tobramycin and to a lesser extent Gen, could promote biofilm formation by *Pseudomonas aeruginosa* through a transcriptional response (19). Based on this observation as well as the possibility that Gen may have more expanded use in the future to treat gonorrheal infections and the relative dearth of knowledge regarding how gonococci respond to sub-lethal levels of clinically used antibiotics, we examined the transcriptional response made by Ng during exposure to sub-lethal concentrations of Gen. Through this work we discovered a Type II toxin-antitoxin (TA) system possessed by Ng and show that its expression can influence biofilm in a strain-specific manner.

## Results and Discussion

### Sub-lethal levels of Gen can induce expression of the *hicAB*-encoded TA system in Ng

We examined whether sub-lethal levels of Gen would induce a transcriptional response in Ng strain FA19 by evaluating growth in GCB broth containing increasing concentrations (0-1X MIC) of Gen. As is shown in Figure S1A, when exposed to Gen at 0.25X MIC (2 μg/mL) Ng grew similarly to the untreated control while higher Gen levels (0.5X and 1.0X MIC) significantly decreased bacterial growth. We found that incubation of strain FA19 with 0.25X of the MIC did not significantly impact biofilm formation compared to the untreated control culture while the higher concentrations (0.5X and 1X the MIC) greatly reduced growth and consequently overall biofilm formation (Fig. S1B).

To determine if a Gen-induced regulon in Ng exists, an RNA-seq analysis was performed using total RNA samples collected from wild-type (WT) strain FA19 grown in the presence or absence of sub-lethal Gen (2 µg/mL) for 4 hours in GCB broth. We identified 23 genes whose level of transcripts changed during Gen exposure (Fig 1A). Of these genes, 21 had increased transcript levels while transcripts from 2 were decreased (Table 1). We sought to validate the RNA-seq data by performing reverse transcriptase-quantitative polymerase chain reaction (RT-qPCR) on transcripts from ten genes. For this purpose, we selected genes randomly to cover the widest linear range for a more accurate correlation analysis. The results showed a correlation coefficient (R²) of 0.9261 indicating similar expression profiles (Fig 1B).

**Fig 1.**
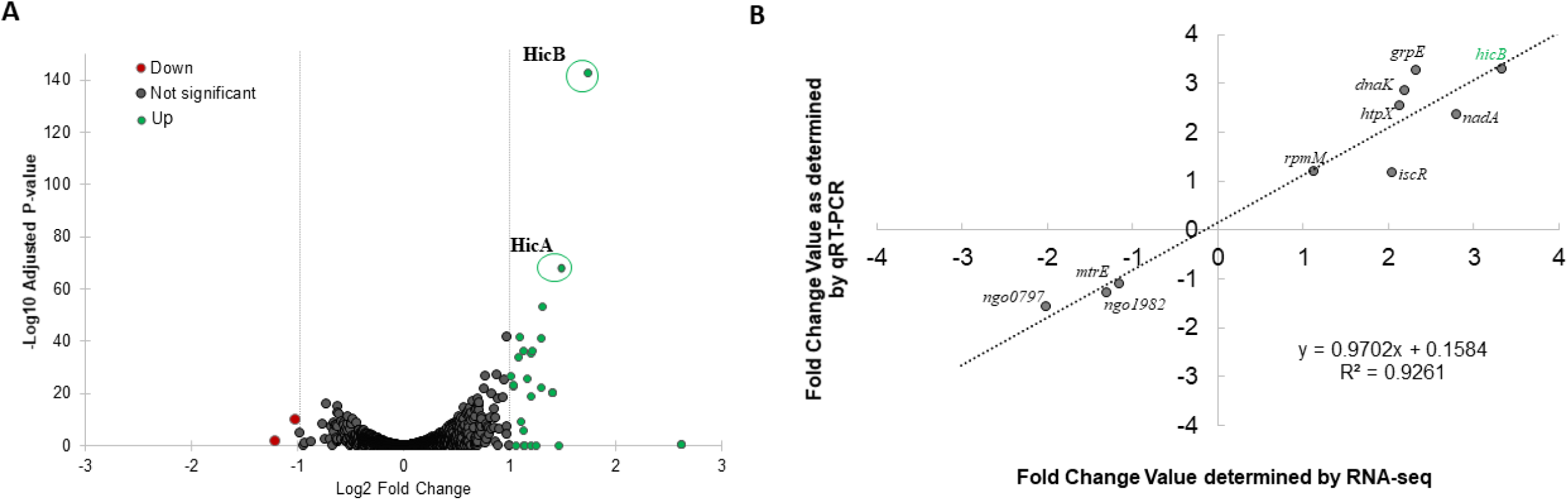
Characterization of the gentamicin induced regulon. A) Volcano plot summarizing differential expression between Ng strain FA19 grown in the presence of sub-lethal gentamicin compared to the absence. Log transformed adjusted p-values were plotted against the log_2_ fold changes for each gene. Genes that are not significantly differentially expressed are shown in gray. Significantly different genes are shown in green (upregulated) or red (downregulated). Location of the *hicA* and *hicB* genes on the plot are circled and labeled. B) Validation of RNA-seq data. RT-qPCR analysis was performed on RNA samples used for RNA-seq as described in methods. Selected genes for analysis are labeled on the plot. The fold change in expression calculated from RT-qPCR analysis (y-axis) was plotted against the value obtained from RNA-seq (x-axis). The correlation between values (R^2^) is shown on the graph.

**Table 1:** Differentially expressed genes in FA19 in the presence of sub-lethal gentamicin compared to the absence.

| Locus_tag<sup>a</sup> | NGO#<sup>b</sup> | Product<sup>c</sup> | Fold Change<sup>d</sup> | P adj<sup>e</sup> |
| --- | --- | --- | --- | --- |
| VT05_RS00725 | NGO_0399 | protease HtpX | 2.14 | 2.45E-42 |
| VT05_RS01940 | NGO_0638 | Fe-S cluster assembly transcriptional regulator IscR | 2.05 | 2.30E-23 |
| VT05_RS02920 | -- | GNAT family N-acetyltransferase | 2.30 | 5.63E-36 |
| VT05_RS02925 | -- | DUF1778 domain-containing protein | 2.02 | 2.83E-27 |
| VT05_RS03080 | NGO_0797 | helix-turn-helix transcriptional regulator | -2.02 | 1.40E-10 |
| VT05_RS04350 | NGO_1046 | ATP-dependent chaperone ClpB | 2.11 | 2.05E-34 |
| VT05_RS04935 | NGO_t22 | tRNA-Ile | 2.65 | 5.71E-21 |
| VT05_RS05470 | NGO_1277 | formylglycine-generating enzyme family protein | 2.47 | 3.06E-23 |
| VT05_RS05625 | NGO_t30 | tRNA-Ile | 2.65 | 5.71E-21 |
| VT05_RS06295 | NGO_1422 | nucleotide exchange factor GrpE | 2.32 | 3.79E-37 |
| VT05_RS06325 | NGO_1428 | hypothetical protein | 2.19 | 1.70E-06 |
| VT05_RS06330 | NGO_1429 | molecular chaperone DnaK | 2.19 | 2.95E-37 |
| VT05_RS06735 | NGO_07995 | pilin-like protein | 2.31 | 1.61E-19 |
| VT05_RS07005 | NGO_1565 | carboxylating nicotinate-nucleotide diphosphorylase | 2.05 | 9.28E-24 |
| VT05_RS07010 | NGO_1566 | hypothetical protein | 2.46 | 1.07E-41 |
| VT05_RS07015 | NGO_1567 | quinolinate synthase NadA | 2.80 | 5.83E-69 |
| VT05_RS07350 | NGO_1627 | type II toxin-antitoxin system HicB family antitoxin | 3.33 | 1.20E-143 |
| VT05_RS07355 | NGO_1628 | type II toxin-antitoxin system HicA family toxin | 2.47 | 8.28E-54 |
| VT05_RS07365 | NGO_1630 | Ortholog of lambda CI repressor | 2.25 | 1.96E-26 |
| VT05_RS07890 | NGO_t39 | tRNA-Ile | 2.65 | 5.71E-21 |
| VT05_RS09065 | NGO_t50 | tRNA-Ile | 2.65 | 5.71E-21 |
| VT05_RS15295 | NGO_RS11400 | IS5/IS1182 family transposase | 2.16 | 5.61E-10 |
| VT05_RS15465 | NGO_RS11480 | IS5/IS1182 family transposase | -2.31 | 0.02839 |

<sup>a</sup>Locus tag as designated by FA19 genome (accession number: NZ\_CP012026)
<sup>b</sup>NGO number as designated in FA1090 genome (accession number: AE004969.1)
<sup>c</sup>Predicted product based on sequence homology or experimental validation
<sup>d</sup>Fold Change: Change in expression in the presence of gentamicin compared to the absence of gentamicin
<sup>e</sup>P adj: Adjusted P-value after correction for multiple comparisons

To better understand the impact of sub-lethal Gen exposure on gene transcript levels, we performed a functional classification analysis using the DAVID tool (20, 21). Functional grouping revealed that most of the genes were classified as related to metabolism and biosynthesis (Table S1). Genes associated with heat shock/protein folding and those of unknown function were the next most represented groups suggesting that exposure of Ng to sub-lethal Gen may induce a stress-like response. Accordingly, we examined the potential overlap of the Gen-induced regulon with other published Ng regulons focusing on stress related regulons. We found that the largest overlap was with the anaerobic regulon with 37% of the genes including *htpX* showing overlap (Table S1) (22). The H_2_O_2_ regulon [27] was the next largest overlap of co-regulated genes at 26.1% which included *grpE, clpB,* and *dnaK.* Taken together, the results indicated that exposure of Ng to sub-lethal Gen can induce expression of stress response genes like those observed for growth of Ng under anoxic conditions or in environments with reactive oxygen conditions.

In addition to the above-mentioned stress response genes harbored by Ng we also noticed that sub-lethal Gen induced expression of two tandemly linked genes NGO1627 and NGO1628, annotated as *hicA* and *hicB*, respectively. These genes are predicted to encode a Type II toxin (HicA)-antitoxin (HicB) system similar to HicAB possessed by other bacteria (Table 1, Fig. 1A); however, in Ng, the genes are in a reverse orientation and predicted to be in a two-gene operon (Fig. S2A) (23). Using RT-qPCR, we verified that a sub-lethal concentration of Gen (0.25X MIC) could induce expression of *hicAB* in strain FA19 (Fig.S2B). During the course of our work the organization of three Ng double-stranded DNA prophages was published with one (Ngo φ3) containing opening reading frames listed as NGO1627, which we assigned as *hicB,* and NGO1630 that encodes an ortholog of the lambda CI repressor (24); the provided information did not assign an open reading frame designated as NGO1628 that we termed as *hicA*

In *E. coli,* HicA is an mRNA interferase that cleaves free mRNAs thus inhibiting translation of selected targets (25). Overexpression of HicA in *E. coli* results in mRNA cleavage and growth inhibition, which can be neutralized by the antitoxin action of HicB. In addition to neutralizing HicA, HicB can serve as a DNA-binding protein and can bind to the operon promoter thereby inhibiting transcription of *hicA* (26, 27). The protein encoded by NGO1628 has 45.2% amino acid similarity (28.6% identity) to the *E. coli* HicA protein (Fig S3, upper panel) but is more similar to the *B. pseudomallei* (74.6% similarity, 62.7% identity) and *Y. pestis* (53.1% similarity; 32.8% identity) HicA proteins. NGO1627, which encodes HicB, only has 43.4% protein similarity (32.1% identity) compared to the *E. coli* HicB, but is also more similar to the *B. pseudomallei* (67.4% similarity, 41.7% identity) and *Y. pestis* (46.9% similarity; 29.2% identity) HicB proteins based on the predicted amino acid sequence structure (Fig S3 lower panel). Of note, while the *E. coli* HicB DNA-binding domain is a classical helix-turn-helix structure, the DNA binding domain of the Ng HicB protein is predicted to be a ribbon-helix-helix (Fig S3 and Fig S4). As TA systems are highly ubiquitous in bacteria, we also compared the HicA and HicB sequences of several representative commensal strains revealing that the proteins in the commensals are highly similar to the Ng proteins (Fig S3). By bioinformatic analysis of a publicly available database (PubMLST) (28) we did not detect the Ng *hicAB* locus in the whole genome sequence *N. meningitidis* strains NMB or MC58. Additionally, we did not detect this Ng sequence in 60 genomes from the *N. meningitidis* urethritis clade (*Nm*UC) isolated from patients with gonorrhea-like urethritis (data not presented) that contain Ng DNA sequences such as the divergently transcribed *aniA-norB* region and the argB, *fHbp*, and *ispD* sequences (29–31). Within the database of these *Nm*UC strains we found 2 additional isolates (isolates #89752242680 and #90135777700) that had an Ng *hicAB* region that was 99% identical to the gonococcal sequence but the two isolates were annotated as being Ng. Thus, unlike other gonococcal genes in these *Nm*UC strains, there no evidence that they contain the Ng *hicAB* locus, but continued surveillance of these isolates is needed to monitor any future appearance.

We next sought evidence that the *hicAB*-encoding proteins in Ng possessed biologic functions like homologous proteins studied in other bacteria. Given the similarity of the Ng HicA to known HicA bacterial toxins, we asked if the Ng HicA could exert a negative impact on bacterial growth. For this purpose, we expressed Ng (strain FA19) *hicA* ectopically from a pET 21a vector in *Escherichia coli*. We found that induction of Ng *hicA* by addition of IPTG could arrest growth of *E. coli*, which was not observed in bacteria containing the vector plasmid alone (Fig. 2A). To determine whether Ng (strain FA19) HicB can function as a DNA-binding protein we assessed its ability to bind the DNA sequence upstream of *hicA* in a specific manner. For this purpose, we performed a competitive electrophoretic mobility shift assay (EMSA). As is shown in Fig. 2B we found that 4 µgs of HicB bound the DNA probe in a specific manner as only an unlabeled specific DNA probe could inhibit such binding. To identify the sequence of the target DNA capable of binding HicB we used a DNase I protection assay. As is shown in Fig. 3, we found that purified HicB can bind to a 14 bp sequence (5ʹ-CAAATACACATAAA-3ʹ) that is positioned 3 bp upstream of a putative −35 hexamer used for *hicAB* transcription. Taken together, we propose that the Ng *hicAB* genes can encode proteins with toxin-antitoxin functions like those identified in other bacteria.

**Fig 2.**
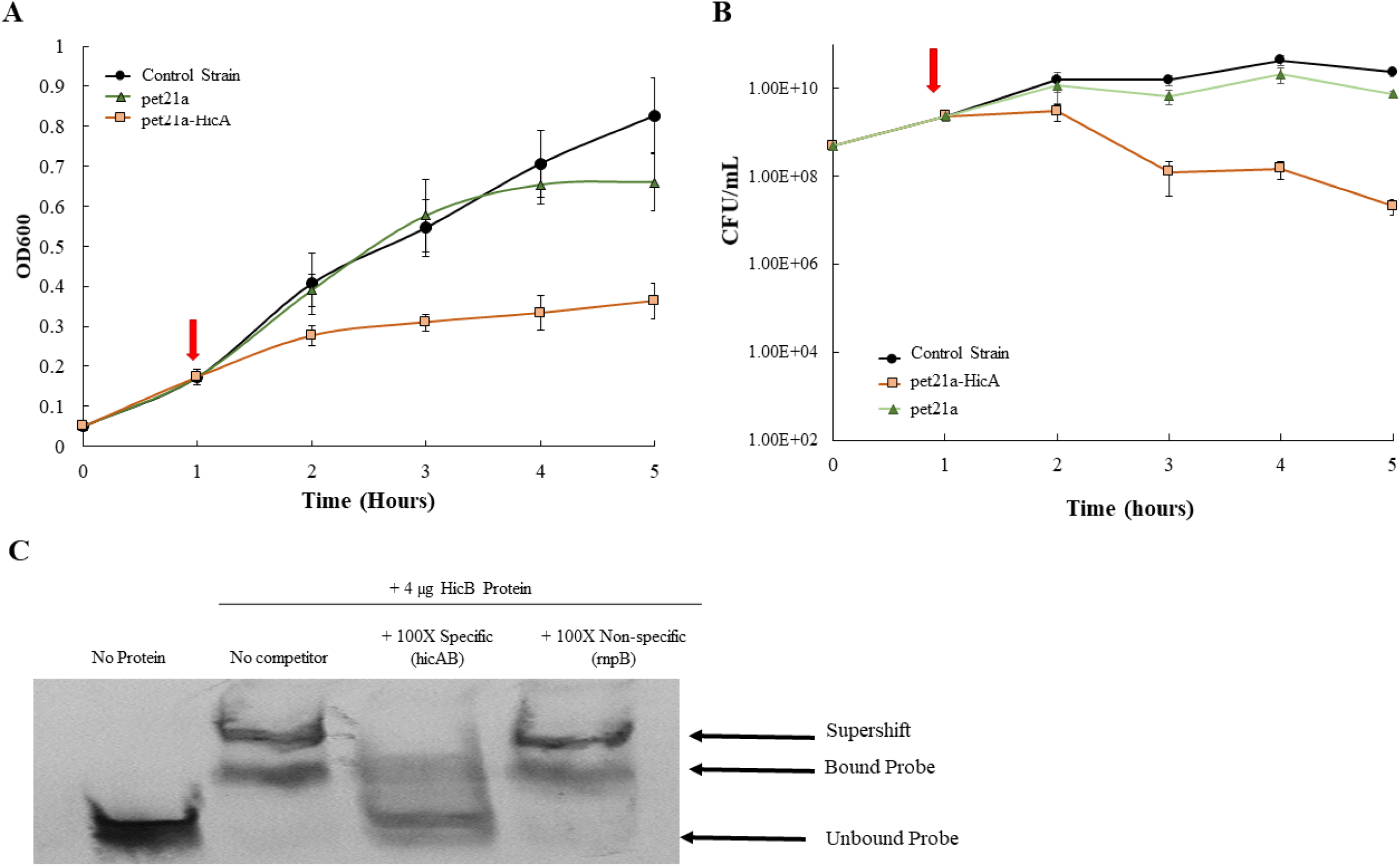
HicA expression is bacteriostatic and HicB binds the *hicAB* promoter. A) HicA expression is bacteriostatic. *E. coli* BL21 strains containing either no plasmid (control), empty vector (pet21a) or HicA under control of IPTG (pet21a-HicA) in the presence of 1 mM IPTG inducer (red arrow). Strains were monitored for 4 additional hours and growth was measured by OD600 (A) and CFU (B). C) HicB binds to the *hicAB* promoter in a specific manner. Shown is a competitive EMSA demonstrating purified HicB binding specificity to the *hicAB* promoter. Lane 1, DIG-labeled probe alone; lane 2 probe plus HicB-His (4 µg); Lanes 3 probe plus 100X concentration of the unlabeled *hicAB* probe (specific); Lanes 4 probe plus 100X concentration of the unlabeled *rnpB* probe (non-specific).

**Figure 3.**
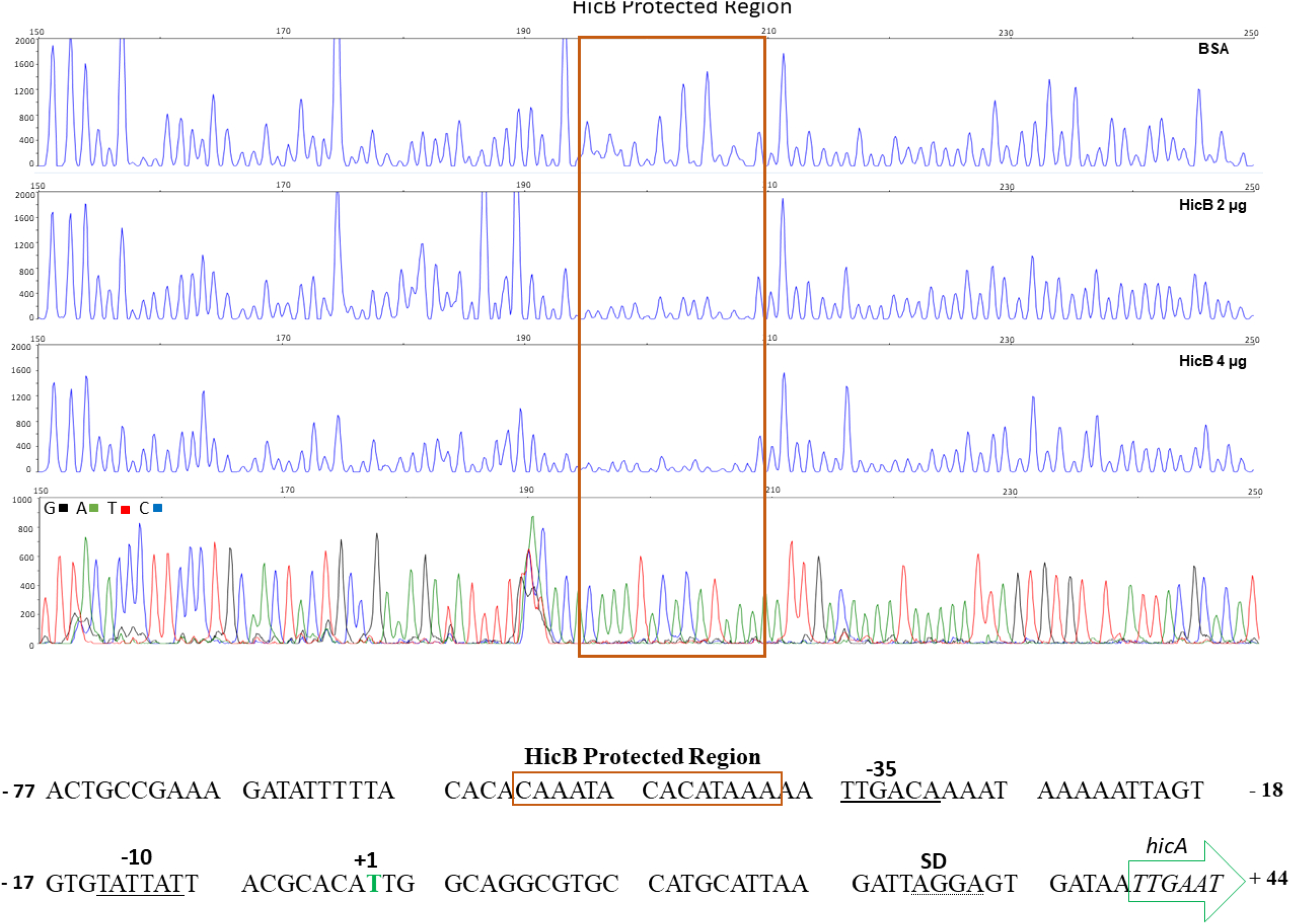
HicB protects a DNA sequence near the *hicAB* promoter. A DNA fragment spanning the *hicAB* promoter region from nucleotide −253 to +49 (relative to the Transcription Start Site, indicated in green) was fluorescently-labeled with 6-FAM (coding strand) and HEX (template strand) and incubated with BSA (control reaction) or HicB prior to digestion with DNase I. The DNase I digestion products were analyzed by capillary electrophoresis. The fluorescence signal corresponding to the 6-FAM probe is shown on the y axis of each electropherogram. Fragment coordinates are shown along the top. The *hicAB* promoter region protected by HicB is boxed. Dideoxy sequencing reactions were manually generated using the 6-FAM primer and a PCR fragment encoding the *hicAB* promoter (bottom panel). DNA sequence of the *hicAB* promoter region is shown below with the HicB protected region boxed.

### Loss of the HicAB TA system can impact the ability of GC to form a biofilm in a strain-specific manner

Earlier studies showed the capacity of Gen to induce biofilm formation by *P*. *aeruginosa* (32), and that loss of *hicAB* reduced the ability of *E. coli* to form a biofilm (33). Based on these observations coupled with our finding that 0.25X Gen could induce expression of *hicAB* (Fig. 1 and Table 1), we evaluated whether loss of these genes would impact biofilm formation by Ng. For this purpose, we constructed *hicAB* null mutants and complemented strains in strain FA19 and two additional strains (FA1090 and F62). The results (Fig. 4A) showed that WT strains FA1090 and F62 expressed statistically significantly more *hicAB* transcripts than WT FA19. Moreover, loss of *hicAB* in strains FA1090 and F62, but not FA19, significantly reduced the ability of Ng to form a biofilm (Figure 4B), which was reversed by complementation with WT *hicAB* expressed ectopically from a pGCC4 construct inserted in the chromosome between *aspC* and *lctP*. Since strains FA19, F62 and FA1090 were isolated from patients over fifty years ago we also tested Ng isolates from the 21^st^ century for their biofilm capability when their *hicAB* region was deleted. For this purpose, we selected strain WHO X (isolated in 2009 from a patient in Kyoto, Japan and the first reported ceftriaxone resistant Ng strain (13) and CDC200 (isolated from a patient in 2012 and resistant to multiple antibiotics (34). As is shown in Fig. S5B the *hicAB::kan* mutants of both WHO X and CDC200 had a reduced biofilm property compared to their parental or complemented strains.

**Fig. 4.**
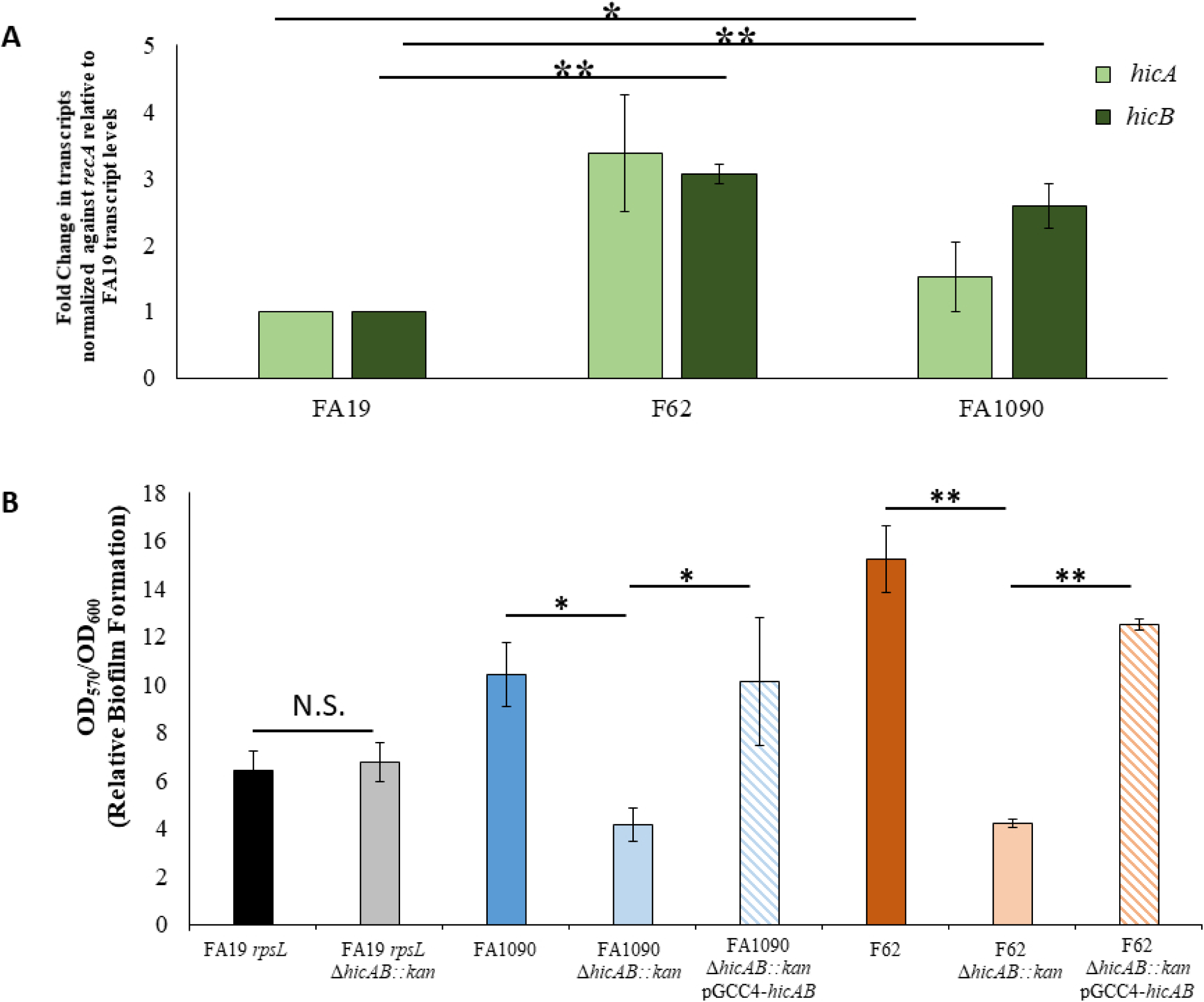
Strain specific differences in *hicAB* expression levels and biofilm formation. A) Levels of *hicA* and *hicB* transcripts in three different *N. gonorrhoeae* genetic backgrounds. RT-qPCR analysis of RNA isolated from NG strains FA19, F62, and FA1090. B) Loss of *hicAB* reduces biofilm formation in FA1090 and F62 strain backgrounds but not FA19. Relative biofilm formation is shown as a ratio of crystal violet stained biomass (OD_570_) corrected by bacterial growth within in each well (OD_600_). Data are representative of at least four independent experiments. Error bars show standard error. *, *P ≤* 0.05; **, *P ≤* 0.01.

Using confocal microscopy, we verified that loss of *hicAB* in strain F62 decreased the ability of Ng to form a biofilm (Fig 5). Importantly, the reduced ability of the F62 *hicAB::kan* mutant to form a biofilm compared to parental strain F62 and the complemented strain was consistent throughout all phases of growth (Fig S5). The reduced ability of strain F62 *hicAB::kan* to form a biofilm compared to parental strain F62, which was not observed in the FA19 background, prompted us to ask if loss of the HicA-HicB TA system in the former would influence other biologic properties of Ng such as susceptibility to antimicrobials and fitness during infection. For this purpose, we examined whether loss of this TA system would alter Ng susceptibility to classical antibiotics, including some used previously or currently in treating gonorrheal infections, and other antimicrobial agents that participate in host defense (a host-derived cationic antimicrobial peptide termed LL-37 _17-32_ and hydrogen peroxide). We found that loss of HicAB did not influence levels of susceptibility of Ng strains FA19 and F62 to any of the examined antibiotics or host antimicrobials (Table S2). Based on the different impact of loss of HicAB on biofilm formation in strain FA19 vs. strain F62, we next sought to investigate the implication of HicAB loss *in vivo* in the dual (WT vs. mutant) experimental lower genital tract female mouse infection model. Consistent with the biofilm data, we found that loss of HicAB did not significantly impact *in vivo* fitness of strain FA19 while such loss significantly reduced *in vivo* fitness of strain F62, as determined by the competitive index, by days 3 and 5 post-infection (Fig. 6).

**Fig 5.**
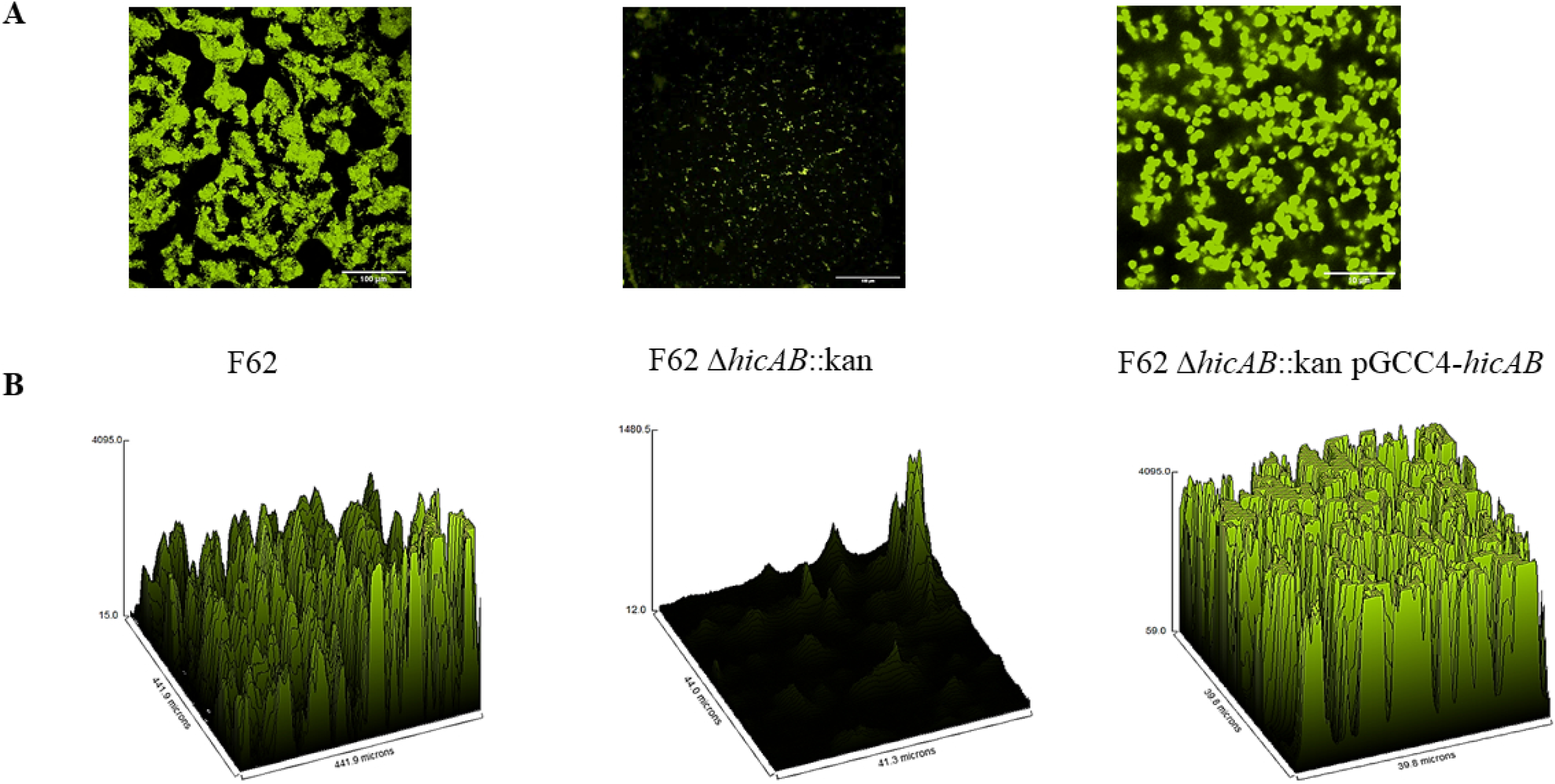
Loss of *hicAB* reduces biofilm formation *in vitro*. A) Impact of *hicAB* deletion on the ability to form biofilms as visualized by confocal microscopy. Tissue culture-treated petri dishes containing glass coverslips were inoculated with strain of *N. gonorrhoeae.* Biofilms were allowed to develop for 24 hours then slides coverslips were stained using the BacLight Bacterial Viability kit to distinguish viable (green) from dead (red) bacteria and analyzed as described within the text. B) Representative 3D surface plot using the maximum-intensity projection of a confocal z-stack from each of the biofilms using x40 magnification with oil emersion. Each plot is a surface of the z-data plotted against x and y.

**Fig. 6.**
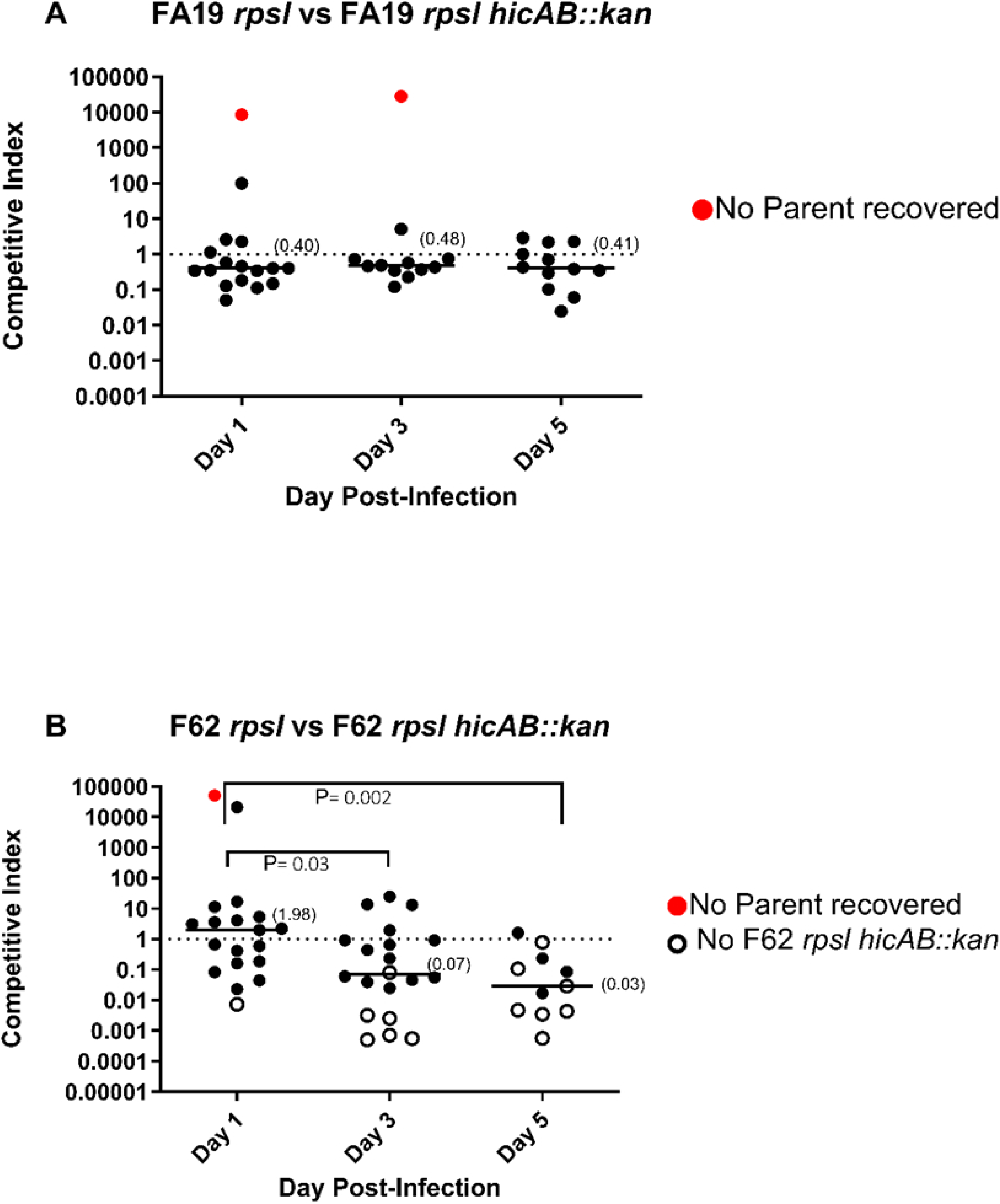
Loss of *hicAB* incurs a fitness disadvantage in a strain-specific manner in an experimental model of genital infection. Groups of female Balb/c mice were inoculated intravaginally with mixed bacterial suspension containing similar CFUs (total dose, 10^6^ CFU *N. gonorrhoeae*; 7 mice/group) of either wild-type FA19 and isogenic FA19 *ΔhicAB::kan* mutant (A) or wild-type F62 *rpsl* and isogenic F62 *rpsl ΔhicAB::kan* (B). Vaginal swabs were collected on days 1, 3, and 5 post-inoculation and suspended in liquid media. Vaginal swab suspensions and inocula were cultured quantitatively on agar with streptomycin (total number of CFUs) and solid media with streptomycin and kanamycin to enumerate *ΔhicAB::kan* mutant strains. Results from two combined experiments are expressed as the competitive index (CI) using the equation CI = [mutant CFU (output)/wild-type CFU (output)]/[mutant CFU (input)/wild-type CFU (input)]. A CI of <1 indicates that the mutant is less fit than the wild-type strain. Red circles indicate a mouse with no wild type bacteria recovered while open symbols designate no mutant CFUs recovered. The limit of detection of 1 CFU was assigned for a strain that was not recovered from an infected mouse. The log10 CI was plotted for each mouse and medians are shown in parentheses. Statistical analysis was performed using Mann-Whitney tests to compare statistical significance of CIs between F62 *rpsl* and isogenic F62 *rpsl* Δ*hicAB::kan*, with *P* values of 0.03 and 0.002 on days 3 and 5 compared to day 1, respectively. Red circles indicate one mouse with no wild type bacteria recovered while open symbols designate no mutant CFUs recovered

### Influence of *hicAB* expression on levels of transcripts by Ng biofilm-associated genes

We questioned whether the reduced biofilm formation observed with the *hicAB* mutant of strain FA1090 was a result of increased death of the mutant bacteria, and if expression of *hicAB* and genes previously linked to biofilm formation was impacted. To this end, we enumerated the percentage of cells within the biofilm well that were in either planktonic (within the supernatant) or biofilm (attached to the surface) states. We found that while 90% of the FA1090 WT strain cells could be found attached to the surface (10% of WT cells were designated planktonic), only 5% of the *hicAB* mutant cells were identified as being in a biofilm with the remaining 95% being found in the planktonic fraction (Fig. 7A); there was no significant difference in the total number of cells within the wells indicating similar growth in our system. Analysis of transcript levels of *hicA* and *hicB* in the biofilm cells compared to the planktonic cells showed that both *hicA* and *hicB* were significantly upregulated in the biofilm; as expected, levels of *hicA* and *hicB* transcripts were both negatively impacted by their deletion (Fig. 7B-C). We also compared transcript levels of *norB, aniA*, and *nuoF* as they were previously shown to be upregulated (*norB, aniA)* and downregulated (*nuoF*) in Ng biofilms formed by strain 1291 (35), and found a similar pattern in WT strain FA1090 (Fig. 7D-F). Interestingly, the *aniA* and *norB* transcript levels appeared to be reduced in the *hicAB* mutant suggesting that this TA system can enhance expression of these genes, which encode enzymes (nitrite reductase and nitric oxide reductase, respectively) that have been shown to be critical for biofilm formation in this Ng strain (35).

**Figure 7.**
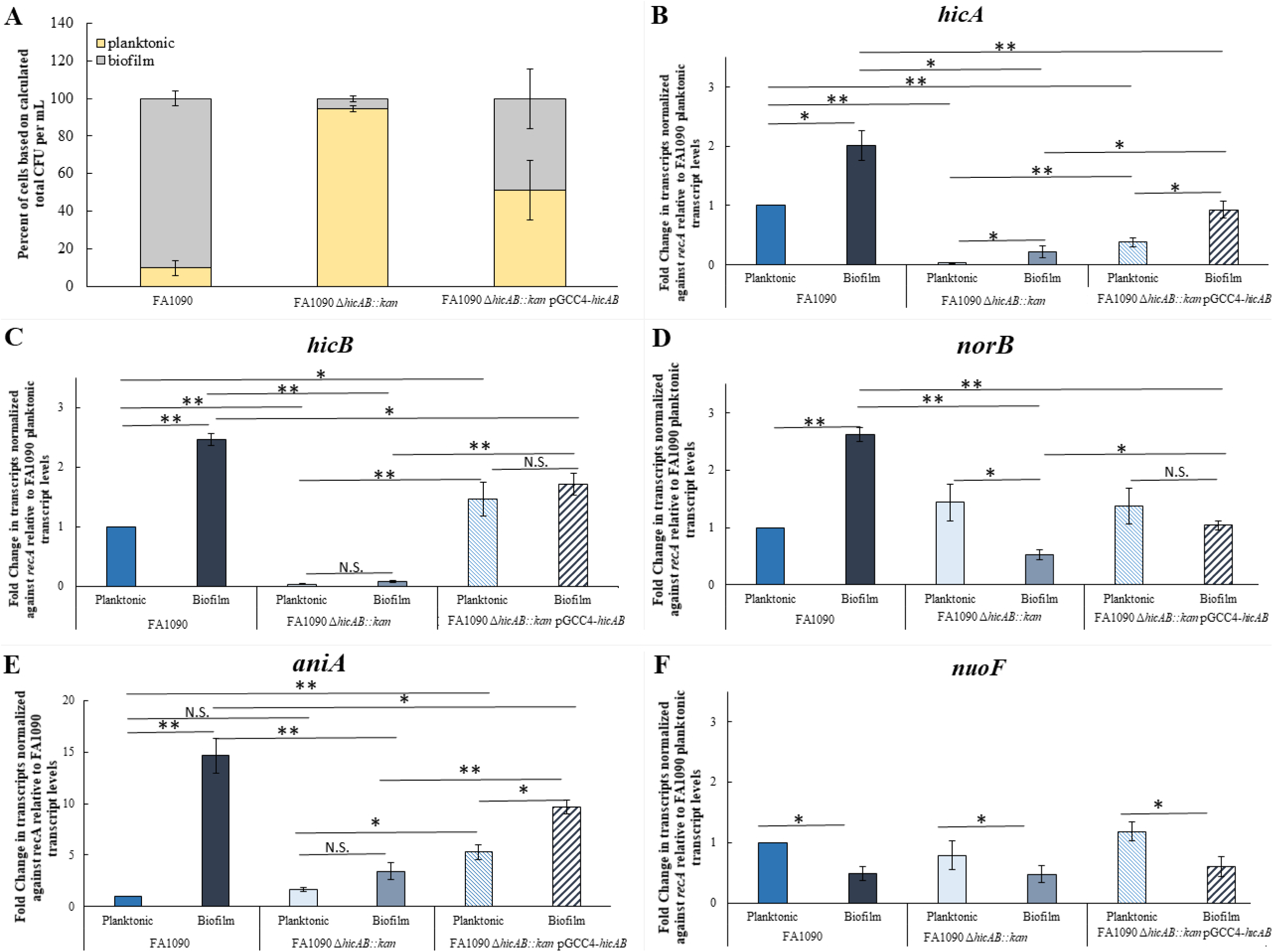
*norB* and *aniA* are differentially expressed in *hicAB* mutant biofilms compared to planktonic cells. A) Colony phenotype of cells in the biofilm assays. Both planktonic (supernatant) and biofilm cells were collected from 96-well plates and counted. Total cell counts were used to calculate the percentage of each phenotype within each strain group and total is set to 100%. Data shown is the average of three independent experiments using colony counts from three technical replicates for each strain and each group (planktonic, biofilm, total). Error bars show standard error. (B-F) Gene expression in biofilm cells compared to planktonic cells. RNA was pooled from biofilm (attached cells) and planktonic cells (supernatant) in triplicate wells and processed for RT-qPCR analysis. Data presented are the fold change of transcripts in the biofilm compared to the planktonic cells (Biofilm/Planktonic) using the WT (FA1090) planktonic value as the comparator (value=1.0). The transcript levels of examined genes were: B) *hicA*; C) *hicB;* D) *norB;* E) *aniA;* F) *nuoF.* Error bars show standard error. *, *P ≤* 0.05; **, *P ≤* 0.01; N.S. = not significant.

### HicAB and the denitrification pathway

The Ng genome contains a partial denitrification pathway encoding AniA and NorB. AniA reduces nitrite to nitric oxide (NO), a chemical that is toxic to Ng in high concentrations (36). To combat this, NorB reduces NO produced by AniA or exogenous NO to the non-toxic nitrous oxide (37, 38). Consistent with the detrimental effect of NO on Ng growth, *aniA* mutants form biofilms with similar overall biomass when compared to the wild-type strain albeit significantly thinner in structure (35). In contrast, *norB* null mutants exhibit severe attenuation in biofilm formation compared to their WT parent (35, 39). Reduced expression of *norB* would therefore limit gonococci’s ability to reduce NO, a condition that could be toxic to Ng. In fact, accumulation of NO is hypothesized to be the root cause of the *norB* mutant’s significantly impaired biofilm formation (35). Given that our *hicAB* mutant biofilms are significantly attenuated in overall biofilm formation, we decided to focus on a potential connection between HicAB and NorB.

Our finding that levels of *norB* and *aniA* transcripts but not that of *nuoF*, was decreased in Ng strain FA1090 *hicAB::kan* compared to parental strain FA1090 along with previous work that the NorB nitrite reductase activity is critical for biofilm formation (35) prompted us to determine if a HicB binding site could be identified in the promoter region between *norB* and *aniA*. We performed a bioinformatic analysis with the FIMO and GLAM2 programs (40, 41) using the HicB binding site identified in the foot printing experiment (Fig. 3) and the *norB/aniA* promoter region. We identified a site upstream of *norB* with a 75% match to the HicB binding site (CAAACAAACTCACATATAGA) (Fig. 8A). To confirm interaction of HicB and this region, we performed an EMSA and found that HicB can bind the promoter region of *norB* (Fig. 8B). This suggests that HicB can potentially directly influence *norB* expression.

**Fig. 8.**
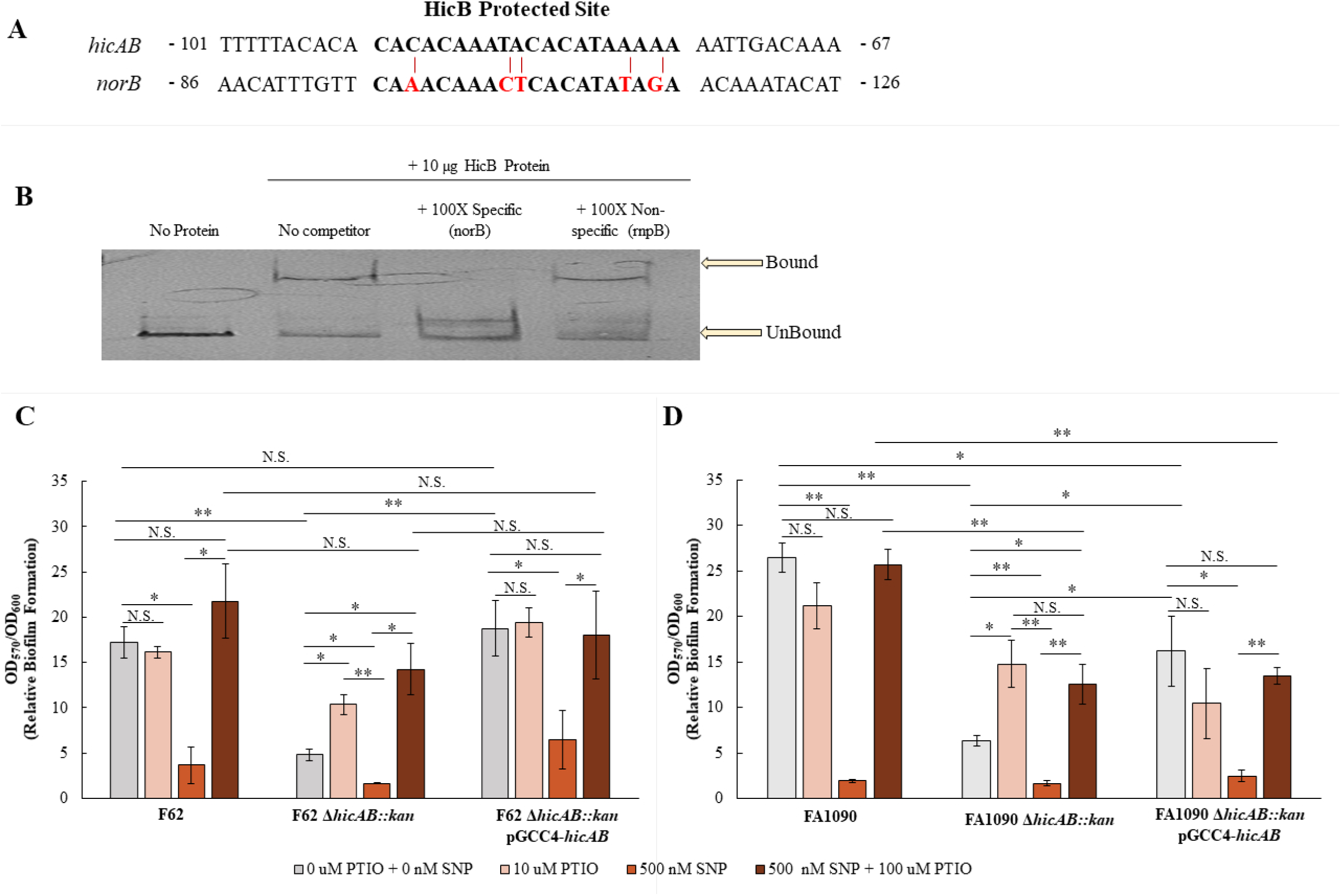
Strain specific differences in *hicAB* transcript levels levels and biofilm formation. A) Alignment of the HicB protected site (bold letters) and the bioinformatically identified region in the *norB* promoter. Numbers represent distance from ATG. Red letters and lines highlight mismatches in the *norB* promoter sequence to the HicB protected site. B) HicB binds to the *norB* promoter in a specific manner. Shown is an EMSA demonstrating purified HicB binding to the *norB* promoter. Lane 1, DIG-labeled probe alone; lane 2 probe plus HicB-His (10 µg); Lanes 3 probe plus 100X concentration of the unlabeled *norB* probe (specific); Lanes 4 probe plus 100X concentration of the unlabeled *rnpB* probe (non-specific). C-D) Addition of NO quencher increases biofilm formation in the *hicAB* F62 (C) and FA1090 (D) backgrounds. Biofilms were either untreated or treated with 10 µM PTIO (NO quencher), 500 nM SNP (NO donor), or simultaneous treatment of 100 µM PTIO and 500 nM SNP. Relative biofilm formation is shown as a ratio of crystal violet stained biomass (OD_570_) corrected by bacterial growth within in each well (OD_600_). Data are representative of at least four independent experiments. Error bars show standard error. *, *P ≤* 0.05; **, *P ≤* 0.01, N.S. = not significant.

Accumulation of NO is toxic to Ng and *norB* mutants are suggested to accumulate NO at concentrations in excess of 500 nM resulting in a significantly impaired biofilm formation phenotype (35). As *norB* expression was decreased in FA1090 *hicAB::kan* compared to parent FA1090, we investigated the effect of NO on biofilms. To determine if excess NO concentrations were responsible for the observed reduction in biofilm formation, we first treated wild-type F62 and its *hicAB* mutant biofilms to increasing concentrations of the NO quencher PTIO (Fig. 8C). We found that 10 µM PTIO significantly increased biofilm formation in the *hicAB* mutant compared to treatment with no PTIO, although it did not restore biofilm formation to WT levels. A similar pattern was found in strain FA1090 and its *hicAB* mutant (Fig.8D). Addition of PTIO did not impact biofilm formation in the WT strains (F62 and FA1090) or in the complemented strains (Fig. 8C-8D). To further examine the effect of excess NO, we next treated biofilms with a sublethal concentration of NO by addition of 500 nM of the nitric oxide donor sodium nitroprusside (SNP), which significantly inhibited biofilm formation in all strains (Fig. 8B-C). Simultaneous treatment of excess NO and NO quencher restored biofilm formation to wild-type levels in all strains. This suggests that excess NO accumulation in the *hicAB* mutant negatively impacts biofilm formation. Interestingly, in the F62 *hicAB* mutant, but not in the FA1090 *hicAB* mutant, the biofilm in the presence of both excess NO and NO quencher, are statistically indistinguishable from the WT F62 strain; this again highlights a strain-specificity in phenotypic outcomes.

## Conclusions

In this work, we determined the Gen-induced regulon following up on our previous studies showing that GC could develop Gen-resistance during growth in the presence of a sub-lethal of this aminoglycoside. We found that when strain FA19 was grown in the presence of 0.25X MIC of Gen that expression of 23 genes was impacted with several associated with a general stress response. Within this set of stress response genes we focused on the two genes encoding proteins similar to the HicAB Type II TA system. We note that the *hicA-hicB* region studied herein is located within the prophage Ngo φ3 region that was recently described (24). Our analysis of the region (designated as NGO1627-1628) indicates it would encode proteins similar to the HicB and HicA, respectively, possessed by other bacteria including many commensal *Neisseria* species. In Ng, the gene order is *hicA* followed by *hicB* as an operon. However, meningococcal strains including NmUC isolates do not seem to harbor the Ng *hicA-hicB* locus described herein.

TA systems in bacteria are ubiquitous gene loci across bacteria and archaea, comprised of a toxin part and its cognate antitoxin part. A typical TA system consists of a stable toxin, which is often a protein and an unstable antitoxin, which can be either an RNA or a protein. In normal conditions, the toxicity of the toxin is neutralized by the antitoxin. Under stress conditions, however, the toxin is released and TA systems play a crucial role in physiology including antibiotic resistance, biofilm and persister formation (42). The *hicAB* operon in *E. coli* is transcribed in response to amino acid and carbon starvation (25, 43). In steady states, the antitoxin and toxin are tightly bound which allows HicB to inhibit HicA activity (26, 44). Within *E. coli,* HicA toxin activity is dependent on Lon protease-mediated degradation of the HicB antitoxin. In fact, most Type II antitoxins in *E. coli* have relatively short half-lives (15-20 minutes) which would allow the cell to rapidly respond to a changing environment (45). This release of HicA would allow RNase action on target mRNA which would temporarily reduce the overall global rate of translation (25). HicA activity induces a reversible cell stasis not death which suggests that activation of the system is meant to be transient not permanent (25, 44). It could therefore be critical for survival under antibiotic stress.

Although much is known about the mechanics of HicA and HicB protein interactions with each other and with their respective targets, the overall biological function of the HicAB, and TA systems in general is still relatively unknown and variations seems to occur across species and niches. In *A. pasteurianus* and *B. pseudomallei,* deletion of HicAB was associated with decreased survival and persister formation, specifically in response to antibiotic stress (46, 47). Incubation with sub-lethal tobramycin, another member of the aminoglycoside family, increased biofilm formation in *P. aeruginosa* on abiotic surfaces (19). Our results indicate that the HicAB TA system is a key factor in response to antibiotic pressure in Ng and we have shown for the first time that an Ng mutant unable to utilize the HicAB TA system for stress response was deficient in biofilm formation. Biofilm formation by Ng is likely an important factor in gonococcal pathogenesis during natural infection (35, 48, 49). HicA may selectively degrade transcripts associated with biofilm formation and natural degradation of the HicB antitoxin under stress conditions may induce a switch from the high-motility state to the low-motility state which would encourage biofilm formation (50–52). Future work to identify potential HicA targets, particularly those associated with biofilm formation, may elucidate HicAB’s involvement in biofilm formation.

In the case of biofilm formation in Ng, we hypothesize that activation of the HicAB system could serve two purposes. HicA mediated RNase activity could result in switching off genes that favor or promote planktonic growth and switching on genes involved in attachment to surfaces. Deletion of *hicAB* in FA1090 and F62 may result in the loss of this switching signal because they do not form a dense biofilm and the majority of the cells remain in the planktonic phase. Inducing a biofilm in Ng could be a resistance mechanism to tolerate an environmental stress without inducing death. Bacteria within the biofilm can exhibit a drastic increase in antibiotic resistance compared to their planktonic counterparts (53). Alternatively, HicA activity may lead to the cell becoming metabolically dormant. HicA overexpression is bacteriostatic not bactericidal (25, 44). In the case of Gen, dormancy can limit its effectiveness because aminoglycosides require active metabolism for action (54). The mechanism and duration of HicA RNase activity has substantial impact on how the Ng responds to stress. Overall, this suggests that the HicAB TA system may have a much more vital role in cell biology and Ng pathogenesis than previously understood.

One of the most interesting findings from this study was the difference in biofilm phenotypes of *hicAB* mutants in different Ng strain backgrounds. Although our Gen regulon work was conducted in FA19, we saw no phenotypic impact on biofilm formation when we deleted the *hicAB* system in the strain. We only saw a phenotype when we deleted *hicAB* in four other Ng strains (FA1090, F62, WHO X and CDC200) suggesting that the property is not uncommon. While it is unclear as to why a difference among strains exists, it was found earlier that *hicAB* was activated by MisR, the response regulator of the MisRS (aka CpxRA) system in the FA1090 background (55), but not in the FA19 background (56), highlighting strain differences in gene expression (57). HicAB may have a more significant biological function in FA1090 and F62 compared to FA19. As the HicAB proteins are identical in the three test strains employed in our work it is likely that other differences such as transcriptional regulatory systems that differentially impact levels of *hicAB* transcripts account for biofilm forming phenotypes associated with this TA system. For example, in FA19 there may exist functional redundancy and/or potential crosstalk with other TA or regulatory systems (45, 58). We have shown previously that the MisR/MisS two-component regulatory system was important for aminoglycoside resistance and that MisR, the response regulator of the system, regulates genes also associated with stress response (56). This study suggests a link between aminoglycoside exposure and stress response in Ng. We propose that this stress response involves increased expression of genes within the Ngo φ3 genetic island. We do not know if the proposed stress response triggers prophage DNA excision and replication or whether transcriptional responses are limited to the genes identified in the Gen regulon without virus production. Lastly, we cannot discount the possibility that the strain-specific differences in biofilm formation due to loss of *hicAB* is linked to Ng strain structural differences of the outer membrane (e.g., outer membrane proteins such as PorB types and/or lipooligosaccharide chemotypess) or secreted proteins that are involved in biofilm formation. Clearly, more extensive investigation with additional Ng strains is needed to understand the influence of the HicA-HicB TA system in Ng biofilm formation.

Denitrification is an important process that allows Ng to thrive in micro-aerobic environments, such as in the urogenital tract, which is critical for its pathogenesis (59, 60). The denitrification pathway in Ng consists of the reduction of nitrite to nitric oxide (NO) via the nitrite reductase AniA and further reduction of NO to nitrous oxide (N_2_O) through nitric oxide reductase NorB (36, 37). Disruption of this pathway could be toxic to Ng. Indeed, expression of an active NO detoxification system has been shown to be important for optimal survival in *Neisseria spp. norB* knockout mutants in Ng exhibit a severe attenuation phenotype in biofilm formation compared to their wild type parent (35, 39). Deletion of *norB* in *N. meningitidis* impairs the bacteria’s ability to colonize and survive within human macrophages and nasopharyngeal mucosal explants (61, 62). Acquisition of the gonococcal *aniA*/*norB* loci was shown to play a key role in the spread and survival of the *N meningitidis* urethritis clade during urethral cell infection (31, 63). All together, these studies show that NorB action is important for pathogenesis in *Neisseria spp* and we observed this as well. Our *hicAB* mutant in F62 exhibits decreased fitness in the mouse model of infection, reduced *norB* expression and reduced biofilm formation which could be partially reversed with NO quencher. Taken together, this suggests that reduced *norB* expression as a result of deletion of *hicAB* could play a significant role in phenotypes exhibited in the *hicAB* mutant. We hypothesize that the elevated levels of *hicAB* transcripts in strains FA1090 and F62 compared to that of FA19 and the ability of HicB to regulate biofilm-associated genes *aniA* and *norB* contribute to the differences in their degree of biofilm formation (Fig. 7). We do not know if this regulatory pathway or differences in post-transcriptional activities (e.g., mRNA decay rates) influence bacterial growth or biofilm formation under anaerobic or microaerophilic conditions. These unknowns will be addressed in our future studies. However, as the amino acid sequences of the HicAB proteins are identical (Fig. S3) in our test strains we do not think the observed differences in target gene transcript levels or biofilm formation are due to alterations in protein function.

To combat growing antibiotic resistance, it is paramount that we understand how Ng responds to environmental pressures and how stress responses influence bacterial susceptibility to antimicrobials. There is potential for HicAB as a novel target for Ng antimicrobial therapy. TA systems have already been under investigation as potential drug targets in other systems (64). The HicAB system could prove a new target for antimicrobial therapy if activation of this system can be shown to significantly influence the biology of Ng or impacts fitness in a mouse model of infection. With growing drug resistance of Ng, targeting a stress response system associated with virulence that also regulates bacterial cell growth could have major impact on controlling the spread of antibiotic-resistant Ng globally. Future work will focus on how HicAB impacts gene expression to influence biofilm formation and pathogenesis *in vivo* as well as how HicAB itself is regulated and activated. Further study of the HicAB system will allow us to better understand its function in Ng biology and antimicrobial resistance.

## Materials and Methods

### Bacterial strains, plasmids, and primers

Ng strains FA19, FA1090, F62 and their isogenic genetic derivative strains, along with the plasmids used and their *Escherichia coli* hosts, are listed in S3 Table. In limited experiments we also used Ng strains WHO X (13) and CDC200 (34). The oligonucleotide primers used in this study are listed in S4 Table. *E. coli* strains were routinely cultured on Luria-Bertani (LB) agar or in LB broth (Difco, Sparks, MD) containing 50 µg/ml kanamycin, 100 μg/ml ampicillin or 100 μg/ml chloramphenicol, as necessary. Gonococci were grown on gonococcal base (GCB) agar (Difco, Sparks, MD) containing Kellogg’s supplements I and II at 37°C under 5.0% (v/v) CO_2_ (65). When strains containing the complementation vector pGCC4 were used, 1 mM IPTG was added to all cultures. Liquid cultures of gonococci for growth assays were begun by inoculating plate-grown cells in pre-warmed GCB broth containing Kellogg’s supplements I and II and 0.043% (w/v) sodium bicarbonate and grown in a 37°C water bath with shaking.

### Growth of bacteria for RNA-seq

Plate-grown FA19 was inoculated into pre-warmed GCB broth containing Kellogg’s supplements I and II and 0.043% (w/v) sodium bicarbonate and grown in a 37°C water bath with shaking for two doublings to ensure cell viability and entry into log phase. The culture was split into two equal volumes and either and either 2 µg/mL of Gen or equivalent volume of sterile vehicle was added to the broth. The cultures were grown for an additional 4 hours then harvested and the pellets stored at −70°C.

### RNA-seq and bioinformatics analysis

RNA-sequencing was performed on the Illumina NextSeq 500 as described by the manufacturer (Illumina Inc., San Diego, CA). Briefly, the quality of the total RNA was assessed using the Agilent 2100 Bioanalyzer. RNA with an RNA Integrity Number (RIN) of 7.0 or above was used for sequencing library preparation. We used the Agilent SureSelect Strand Specific mRNA library kit as per the manufacturer’s instructions (Agilent, Santa Clara, CA). Library construction began with ribosome reduction using the RiboMinus rRNA depletion kit for bacteria as described by the manufacturer (Invitrogen). The resulting RNA was randomly fragmented with cations and heat, which was followed by first strand synthesis using random primers with inclusion of Actinomycin D (2.4ng/µL final concentration). Second strand cDNA production was done with standard techniques, the ends of the resulting cDNA were made blunt, A-tailed and adapters ligated for amplification and indexing to allow for multiplexing during sequencing. The cDNA libraries were quantitated using qPCR in a Roche LightCycler 480 with the Kapa Biosystems kit for Illumina library quantitation (Kapa Biosystems, Woburn, MA) prior to cluster generation. Cluster generation was performed according to the manufacturer’s recommendations for onboard clustering (Illumina). Raw read statistics are available in Table S5.

For the bioinformatics analysis, fastq sequences were deposited to the public Galaxy tool shed (66). Briefly, paired-end fastq files generated by the sequencing platform were aligned to the *N*. *gonorrhoeae* FA19 genome (GenBank assembly accession: GCA_000273665.1) using the Bowtie2 tool to generate BAM files. Counts were calculated from the bam files using the htseq-count tool and differential expression between samples determined using the DEseq2 tool. The fastq and BAM files are available in the Gene Expression Omnibus (GEO) (67) with GEO series accession number GSE291660.

### Construction of *hicAB* mutants and complementation

In order to ensure complete removal of the *hicAB* operon, a Δ*hicAB::*kan mutant was constructed as follows. Primer pairs hicABstart/hicAB For and hicABstop/hicAB Rev were used to amplify the upstream and downstream sequences of the *hicAB* gene. The two PCR products were ligated together at the XbaI site and cloned into pBad TOPO-TA. The confirmed plasmid was designated pBad-*hicAB*. pBad-*hicAB* was digested with HindIII and the ends blunted with Klenow. The *aphA-3* non-polar Kan^R^ cassette was amplified from pUC18K using PFU to create blunt ends. The Kan^R^ cassette was ligated to pBad-*hicAB* to create pBad-*hicAB*::kan. This plasmid was used to transform FA1090 and F62. Deletion of the gene and loss of the cassette was confirmed by PCR and sequencing. The resulting strains were designated FA1090 Δ*hicAB::*kan and F62 Δ*hicAB::*kan. Complementation was performed as previously described using the pGCC4 complementation vector (68). Briefly, the entire *hicAB* operon was amplified from FA1090 genomic DNA using primers hicABpacI and hicABpmeI and inserted into digested pGCC4 vector. The resulting plasmid was used to transform FA1090 Δ*hicAB::*kan and F62 Δ*hicAB::*kan and colonies were selected on erythromycin 2 µg/mL Transformants were verified by PCR and sequencing using primers lctp and aspC1. To make the strains Str^R^, the *rpsL* allele from previously described FA19 *rpsL* (69) was cloned into pBad and used to transform the F62 and F62 Δ*hicAB::*kan. Strains were selected on streptomycin 100 µg/mL and confirmed by Sanger sequencing of PCR-amplified *rpsL* gene products.

### Analysis of Ng transcript levels

Gonococci were harvested at late-log phase and the pellets were stored at −70°C. For validation of RNA-seq, an aliquot of the RNA used for RNA-seq was utilized. All RNA was purified by Trizol extraction as per manufacturer instructions (Thermo Fisher Scientific, Waltham, MA, USA) followed by Turbo DNA-free (Ambion, Austin, TX, USA) treatment. cDNA was generated using a QuantiTect reverse transcriptase kit (Qiagen, Venlo, The Netherlands). RT-qPCR was conducted using the IQ SYBR Green Supermix and a CFX Connect Real Time System (Bio-Rad Laboratories). We validated our RT-qPCR methods by examining primer efficiency, primer specificity (melt temperature), and linear dynamic range for each primer pair utilized herein. For RT-qPCR analysis, relative expression values were calculated as 2^(CT^ ^reference−^ *^CT^* ^target)^, where CT is the fractional threshold cycle (70). The level of housekeeping reference gene *recA* mRNA was used as internal reference. All RT-qPCRs were performed in technical duplicates and biological triplicates.

### Minimum inhibitory concentration (MIC) determination

The MICs of antibiotics were determined by the agar dilution method (71). Antimicrobials were purchased from Sigma Chemical Co. (St. Louis, MO). Plates were incubated for 48 hours at 37°C under 5.0% (v/v) CO_2_-enriched atmosphere before being analyzed. The susceptibility of Ng strains to hydrogen peroxide and LL-37_17-32_ (kindly provided by E. Yu, Case Western University, Cleveland, Ohio) were determined as described previously (72, 73).

### Analyses of *N. gonorrhoeae* biofilms

5×10^5^ of each gonococcal strain was seeded into 96 well microtiter plates (Costar, Corning, New York) and incubated at 37°C for 24h to allow static biofilm formation. Bacterial density was then read (absorbance at 600 nm) using a Victorx3 2030 Multilabel Reader (Perkin Elmer, Waltham, MA) to determine the baseline growth for each well. After incubation, planktonic bacteria were removed, and the wells were gently washed 3 times with phosphate-buffered saline. Plates were allowed to dry and then stained for 30 mins with 0.1% (w/v) crystal violet(CV). Plates were washed thoroughly 3 times with phosphate-buffered saline then dried. CV-stained biofilms were solubilized with DMSO and absorbance, indicative of biofilm biomass, was read at 570 nm. Data were adjusted for background and each assay was performed using quadruplicate wells in at least 3 independent experiments. Relative biofilm formation was calculated as ratio of the OD_570_/ OD_600_ for each well and averaged across strains. To determine colony type (planktonic or biofilm), cells were seeded into 96-well plates as described above. For planktonic cells, the supernatant was carefully removed from the well so as not to disturb the biofilm layer and cells were serially diluted, and CFU/mL determined. Biofilm cells were harvested by suspending the biomass attached to the well bottom in 100 µl GC broth and gently washed by pipetting up and down until no biofilm was visible in the well bottom. Total colony counts were performed by removing the supernatant and attached cells in the same suspension. The total cell count was used to calculate the percentage of each cell type [e.g. (# planktonic cells/ mL) / (# total cells per mL)]. Total cell counts calculated from the sum of planktonic and biofilm were compared to total cell counts calculated. Data is the average of three independent experiments using colony counts from three technical replicates for each strain and each group (planktonic, biofilm, total). For analysis of gene expression in biofilm cells compared to planktonic cells, RNA was pooled from biofilm (attached cells) and planktonic cells (supernatant) in triplicate wells and processed for qRT-PCR analysis as described above. Planktonic cell supernatants were collected, pelleted, and stored at −70°C. For biofilm cells, 200 µl GC Broth was added to each well and attached cells were removed until each well was visually clear. Collected cells were then pelleted and stored at −70°C until processing.

### Confocal Microscopy of Biofilms

For confocal microscopy of gonococcal biofilms on an abiotic surface, 1 × 10^6^ of each strain was inoculated onto 35 mm × 10 mm petri dishes (Falcon, Corning, New York) containing glass coverslips. Samples were incubated (37°C, 5% CO_2_) for 24 hours to allow biofilm formation. Biofilms were stained using the Live/Dead BacLight Bacterial Viability kit (Thermo Fisher), according to the manufacturer’s instructions. Analyses of the biofilms was achieved using a Nikon A1R HD25 (Nikon Instruments Inc., Melville, NY) laser scanning confocal microscope. Images were captured using diode laser wavelengths of 488 nm (live bacteria) and 561 nm (dead bacteria) and collected as 1-μm Z-stack slices under 40× oil emersion. Biofilm architecture (surface plots) was examined using FIJI software (74).

### Biofilm treatment with nitric oxide quencher and donor

For determining the impact of nitric oxide on biofilm formation in the *hicAB* mutant, biofilm assays were performed as previously described with the following additions. Biofilms were treated with 10 µM nitric oxide quencher 2-phenyl-4,4,5,5-tetramethylimidazoline-1-oxyl 3-oxide (PTIO) at the start of the biofilm assay. The biofilms were allowed to develop in the 96 well plate for 22-24 hours at 37°C with 5% CO_2_. As a control, excess NO provided by 500 nM nitric oxide donor sodium nitroprusside (SNP) was utilized. To demonstrate that the effects of SNP were due to nitric oxide release, in another control, 100 µM PTIO was added simultaneously with 500 nM SNP. Untreated wells were grown at the same time. All samples were run in quadruplicate wells per strain in four independent biological replicates.

### DNase I Protection Assay

A DNA fragment spanning the *hicAB* promoter region from nucleotide −253 to +49 (relative to the Transcription Start Site) was fluorescently-labeled with 6-carboxyfluorescein (FAM)- and 6-carboxy-2’,4,4’,5’,7,7’-hexachlorofluorescein (HEX)-labeled primers hicAB_FP_For and hicAB_FP_Rev. DNase I digestion reactions using purified HicB protein were performed as described previously (75). Detection of the DNase I digestion peaks was conducted in a 3730 capillary sequencer (Applied Biosystems) and the alignment of the corresponding electropherograms was generated using Peak Scanner software(Applied Biosystems). A negative control reaction was conducted using 4 µgs of BSA. Dideoxy sequencing reactions were manually generated using the 6-FAM primer and a PCR fragment encoding the *hicAB* promoter.

### Electrophoresis Mobility Shift Assay (EMSA)

EMSAs were conducted using the second-generation digoxigenin (DIG) gel shift kit (Roche Applied Sciences, Madison, WI) as previously described (72). DNA fragments were generated by PCR from *N*. *gonorrhoeae* FA1090 gDNA with primer pairs hicAB_P1_For / hicAB_P1_Rev for the *hicAB* probe, norB EMSA F/norB EMSA R for the *norB* probe and rnpB1/rnpB2 for *rnpB* probe. *hicAB* and *norB* probe was labeled with digoxogenin (DIG) following manufacturer’s protocol. Specificity of HicB binding to the DIG-labeled *hicAB* and *norB* promoters was assessed by adding 100-fold excess of unlabeled DNA fragments encoding either the specific competitor or a non-specific DNA competitor (*rnpB)* purified HicB protein for 30 min at 30°C in Binding Buffer: 20 μL of 20 mM Hepes, pH 7.6, 1 mM EDTA, 10 mM (NH_4_)_2_SO_4_, 1 mM DTT, 0.2% (v/v) Tween-20, 30 mM KCl and 1.25 ng/μL Type XV calf thymus DNA. Reactions were separated by electrophoresis in 5% Mini-Protean TBE Precast Gels (Bio-Rad) and transferred to nylon membranes and crosslinked. Blots were developed using an anti-DIG Fab fragment-AP conjugate.

### Competitive female mouse infection model

Competitive infections were conducted as described (76). Female BALB/c mice (6 to 8 weeks old; Charles River Laboratories Inc., Wilmington, MA; NCI Frederick strain of inbred BALB/cAnNCr mice, strain code 555) were treated with 0.5 mg of 17β-estradiol given two days prior to, the day of, and two days after bacterial inoculation to increase susceptibility to *N*. *gonorrhoeae*. Mice were also given antibiotics to suppress the overgrowth of commensal bacteria that occurs under the influence of estrogen. Groups of mice were inoculated vaginally with similar numbers of FA19 and isogenic FA19 *hicAB::*kan bacteria (total dose 10^6^ CFU; 8 mice/group) or F62 and isogenic F62 *hicAB::*kan (total dose 10^6^ CFU; 7-12 mice/group). Vaginal swabs were collected on days 1, 3, and 5 post-inoculation and suspended in 1mL GCB. Swab suspensions and inocula were cultured quantitatively on GCB supplemented with Str (FA19 or F62; total number of CFUs) and GCB with Str and Kan (FA19 *hicAB::kan* or F62 *hicAB::kan*; CFU). Results are expressed as the competitive index (CI) using the equation CI = [mutant CFU (output)/wild-type CFU (output)]/ [mutant CFU (input)/wild-type CFU (input)]. The limit of detection of 1 CFU was assigned for a strain that was not recovered from an infected mouse. A CI of >1 indicates that the mutant is more fit than the WT strain. Statistical analysis was performed using Mann-Whitney tests.

### Bioinformatic analysis of bacterial whole genome sequences

Analysis of bacterial whole genome sequences for similarity to the Ng *hicA-hicB* region of strain FA19 was conducted as described by Holley et al. (9) using publically available sequences in the PubMLST database (28).

### Statistical methods

All data except animal infection model data are expressed as means with standard error (SE). Statistical significance between all quantitative data was analyzed by Student t-tests or ANOVA followed by Tukey’s honestly significant difference post-hoc test. Statistical significance was set at *P* < 0.05.

### Ethics Statement

Animal experiments were conducted at the Uniformed Services University of the Health Sciences (USUHS) according to the guidelines of the Association for the Assessment and Accreditation of Laboratory Animal Care under a protocol approved by the University’s Institutional Animal Care and Use Committee. The USUHS animal facilities meet the housing service and surgical standards set forth in the “Guide for the Care and Use of Laboratory Animals” NIH Publication No. 85–23, and the USU Instruction No. 3203, “Use and Care of Laboratory Animals”. Animals are maintained under the supervision of a full-time veterinarian. For all experiments, mice were humanely euthanized by trained personnel upon reaching the study endpoint using a compressed CO_2_ gas cylinder in LAM as per the Uniformed Services University (USU) euthanasia guidelines (IACUC policy 13), which follow those established by the 2020 American Veterinary Medical Association Panel on Euthanasia (https://www.usuhs.edu/mps/facilities-resources).

## Acknowledgments

We thank the University of Alabama-Birmingham Genomics and Bioinformatics Cores for RNA-seq and preliminary regulon analysis. We also thank the Emory Integrated Cellular Imaging Core for assistance with confocal microscopy.

The contents of this article are solely the responsibility of the authors and do not necessarily reflect the official views of the National Institutes of Health, the U.S. Department of Veterans Affairs, the U.S. Department of Defense, or the United States government.

We thank E.W. Yu for providing antimicrobial peptide LL-37_17-32_.

This work was supported by NIH grants R01 AI147609 and R01 AI021150 (W.M.S.). W.M.S. is the recipient of a Senior Research Career Scientist Award from the Biomedical Laboratory Research and Development Service of the U.S. Department of Veterans Affairs. Confocal Microscopy was supported in part by the Integrated Cellular Imaging shared resource of Winship Cancer Institute of Emory University and NIH/NCI under award number P30CA138292. The funders had no role in study design, data collection and analysis, decision to publish, or preparation of the manuscript. The views expressed in this manuscript are those of the authors and do not reflect those of the National Institutes of Health, the Department of Defense, or the Department of Veterans Affairs.

## Data availability Statement

Data underlying the transcriptome analysis are available at Gene Expression Omnibus (GEO) under ID #: GSE291660. All other relevant data are in the manuscript and its supporting information files.

